## Supplementary material for "Antifungal alternation can be beneficial for durability but at the cost of generalist resistance": Sipplementary Material

**Supporting Information Figure S1:** **Variation of resistance evolution in selection regimes**

ρ, the mean resistance evolution rate, is shown for each selection regime. The error bars represent the standard error *P*-values for pairwise comparisons were obtained with linear models (Tukey’s *post-hoc* correction). Alternations involving B, C or P had a beneficial or neutral effect on resistance evolution rate relative to the continuous use of the same fungicide. The benefit of alternation depends on the intrinsic resistance risks of the paired AIs.

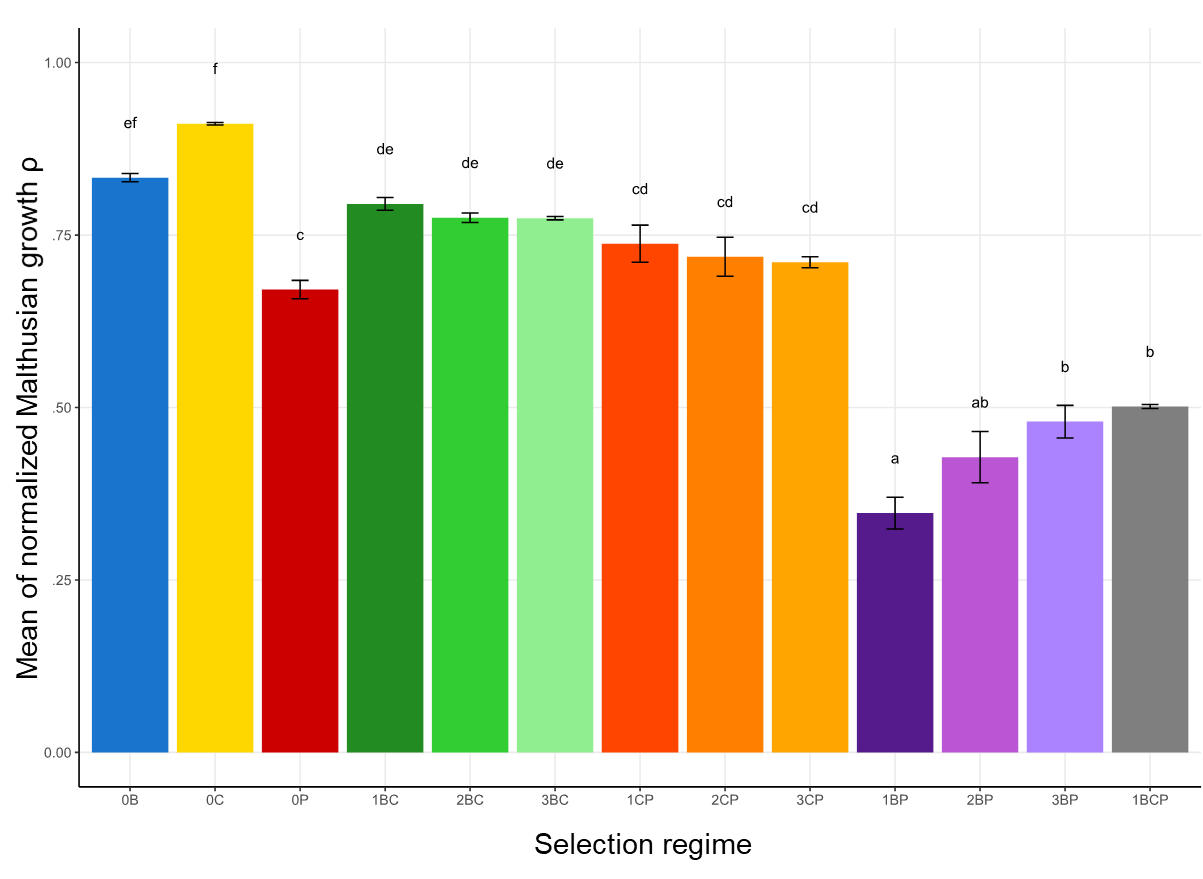

**Supporting Information Figure S2:** **Expression of *tub2*, *sdhC*, *cyp51* and *mfs1* in evolved strains of *Z. tritici***

Relative gene expression (expressed as 2^-ΔΔCt^) of *tub2*, *sdhC*, *cyp51*, encoding the target proteins of carbendazim, benzovindiflupyr and prothioconazole-desthio, respectively, shown for eight isolates collected from various selection regimes. The expression of the drug transporter MFS1 was also quantified, as enhanced efflux is recognized as a generalist resistance mechanism in field isolates of *Z. tritici.* Isolates were chosen to represent the diversity of sensitivity the various fungicides, as shown from their scores in droplet tests (shades of brown, on the top of each graph). The ancestral strain IPO-323 was added as negative control. Two biological replicates were tested independently. The expression of *tub2*, *cyp51* and *mfs1* was normalized to that of the genes encoding EF1α (black bars) and ubiquitin (light grey bars). The expression of *sdhC* was normalized to that of the gene encoding β-tubulin (grey bars).

The overexpression of the three target genes was not observed among isolates resistant to B, C or P. The overexpression of *msf1* is noticed for few isolates exhibiting moderate to high resistance to tolnaftate, suggesting that other unknown additional mechanisms must be involved.

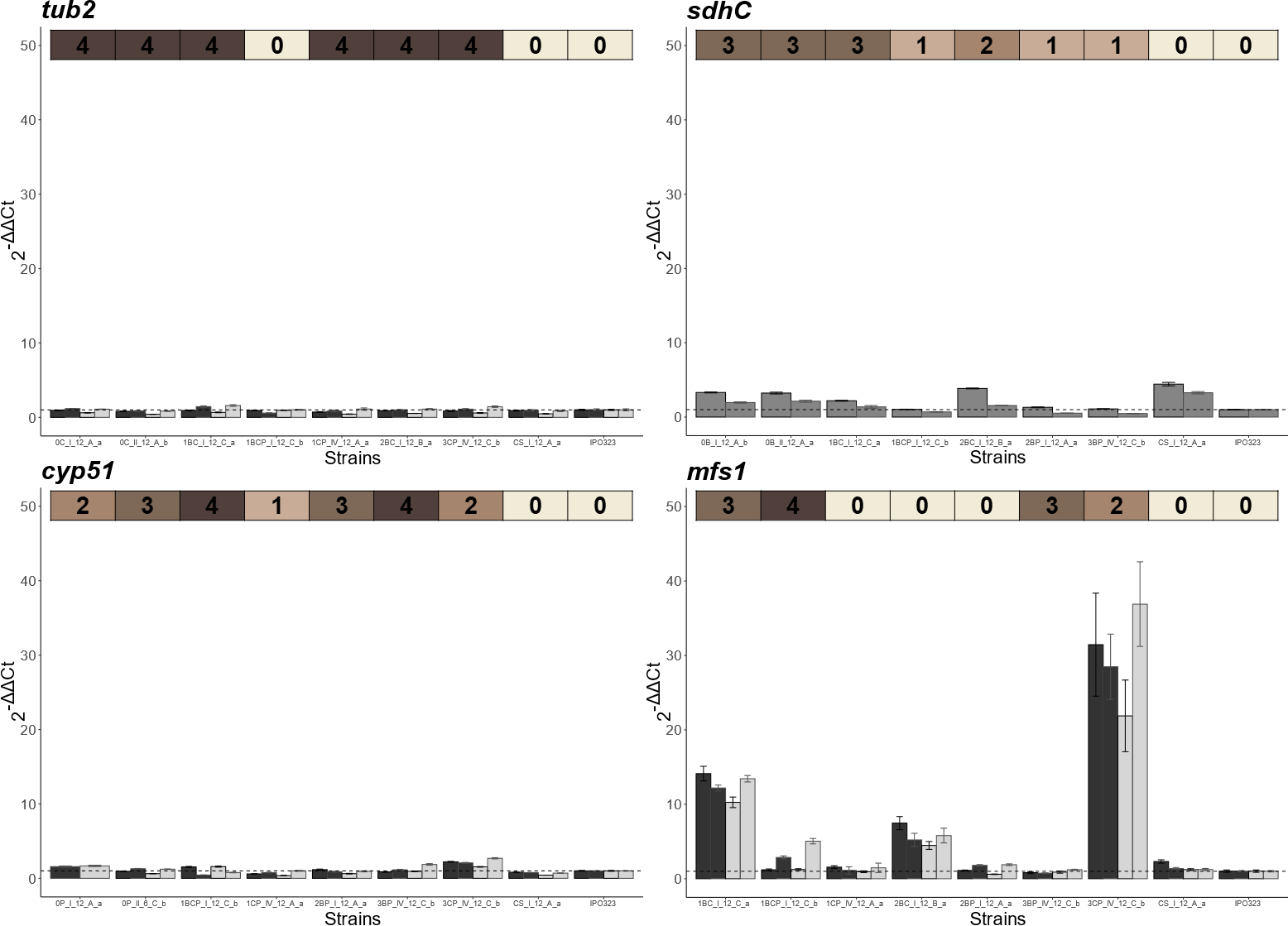

**Supporting Information Methods S1:** **Statistical programs used for data analysis**

All analyses were performed and all figures were generated with R 4.0.4 and R Studio 1.4.1106.

R Core Team. *R: A Language and Environment for Statistical Computing*. (R Foundation for Statistical Computing, 2021).

RStudio Team. *RStudio: Integrated Development Environment for R*. (RStudio, PBC, 2021).

The additional packages used for this work are listed below.

- **car**: Fox, J. & Weisberg, S. *An R Companion to Applied Regression*. (Sage, 2019).
- **cowplot:** Wilke, C. O. *cowplot: Streamlined Plot Theme and Plot Annotations for ‘ggplot2’*. (2020).
- **coxed:** Kropko, J. & Harden, J. J. *coxed: Duration-Based Quantities of Interest for the Cox Proportional Hazards Model*. (2020).
- **emmeans**: Lenth, R. V. *emmeans: Estimated Marginal Means, aka Least-Squares Means*. (2021).
- **FactoMineR**: Lê, S., Josse, J. & Husson, F. FactoMineR: A Package for Multivariate Analysis. *Journal of Statistical Software* **25**, 1–18 (2008).
- **ggplot2:** Wickham, H. *ggplot2: Elegant Graphics for Data Analysis*. (Springer-Verlag New York, 2016).
- **ggpubr:** Kassambara, A. *ggpubr: ‘ggplot2’ Based Publication Ready Plots*. (2020).
- **ggdendro:** Vries, A. de & Ripley, B. D. *ggdendro: Create Dendrograms and Tree Diagrams Using ‘ggplot2’*. (2020).
- **Multcomp:** Hothorn, T., Bretz, F. & Westfall, P. Simultaneous Inference in General Parametric Models. *Biometrical Journal* **50**, 346–363 (2008).
- **nlme:** Pinheiro, J., Bates, D., DebRoy, S., Sarkar, D., & R Core Team. *nlme: Linear and Nonlinear Mixed Effects Models*. (2021).
- **survival**: Therneau, T. M. *A Package for Survival Analysis in R*. (2020).

**Supporting Information Methods S2:** **Sequencing fungicide target genes in *Z. tritici***

1. **Primers used to sequence fungicide target genes and to detect alterations of the *MFS1* promoter in *Z. tritici***

| **Primer name** | **Primer sequence** | **Tm** (°C) | **Reference** |
| --- | --- | --- | --- |
| ***Tub2*** | | | |
| Tub5 | CAGTCACTTGCAAACGGGTA (F) | 63.7 | This work |
| Tub14 | GAAGCCTCCTGGTACTGCTG (R) | 63.9 | - |
| Tub15 | GTCGACCAGGTTCTCGATGT (F) | 64.1 | - |
| Tub13 | AACCGACCATGAAGAAGTGG (R) | 63.8 | - |
| ***cyp51*** | | | |
| CypFra2r | TCCTTCTCCTCCCTCCTCTC (R) | 64.0 | Brunner *et al* (2008) |
| EBI112 | TTCAGCACGCTCGCCATCCTCC (F) | 76.5 | Brunner *et al* (2008) |
| Cyp9 | TCTATACTGCCAGCCGATCA (R) | 63.3 | This work |
| Cyp12 | CACTAACAGGCCAACTGCAA (F) | 63.8 | - |
| NB1 | CACTCTTCATCTGCGACCGAGTC (R) | 69.4 | Leroux *et al* (2007) |
| ***sdhB*** | | | |
| Mg SDHB | TTCGGCACGTCGAGTTTCTGGCCATGCTGGCGG (F) | 89.2 | This work |
| Mg SDHB Rv | CCACAATGCCGCAAAATCGGCACAACTGCTCCC (R) | 86.1 | - |
| Mg SeqSDHB Fw | ACATCTACCGATGGAACCC (F) | 61.2 | - |
| Mg SeqSDHB Rv | TGTGGCATCGGTACAAGC (R) | 63.5 | - |
| ***sdhC*** | | | |
| Mg SDHC Fw \|; Mg SDHC Rv \| | TCTCTTCGTCGACATCACCA (F)  ACGCACTCGCAACACTCA (R) | 64.7  64.7 | This work  - |
| ***sdhD*** |  |  |  |
| Mg SDHD Fw | ACAGCTTCCGAGGTTTCGCG (F) | 70.8 | This work |
| Mg SDHD Rv | GCCATCTGTATATACACGCCC (R) | 63.0 | - |

Brunner, P. C., et al. (2008). "Evolution of the CYP51 gene in *Mycosphaerella graminicola*: evidence for intragenic recombination and selective replacement." Molecular Plant Pathology **9**(3): 305-316.

Leroux, P., et al. (2007). "Mutations in the CYP51 gene correlated with changes in sensitivity to sterol 14 alpha-demethylation inhibitors in field isolates of *Mycosphaerella graminicola*." Pest Management Science **63**(7): 688-698.

2. **Regular PCR Mix**

| Mix | 1 tube (µL) |
| --- | --- |
| DNA | 1 |
| dNTP (25mM) | 0.5 |
| FW (10 µM) | 1 |
| RV (10 µM) | 1 |
| Titanium buffer (10X) | 5 |
| Titanium *Taq* polymerase (50X) | 0.5 |
| Deionized H_2_O | 41 |
| Total volume | 50 |

DNA polymerase: Titanium Taq (Takara Bio)

3. **Regular PCR program**

| Initiation | 95°C | 3 min |  |
| --- | --- | --- | --- |
| Denaturation | 95°C | 30 s |  |
| Annealing | Tm°C | 30 s | 35 cycles |
| Elongation | 72°C | 1min |  |
| Final | 72°C | 5 min |  |
|  | 10°C | To the end of the cycle |  |

**Supporting Information Methods S3:** **Detecting polymorphism in promoters of target genes in *Z. tritici***

1. **Primers used to detect length polymorphism in the promotor sequence of fungicide target genes in *Z. tritici***

| **Primer name** | **Primer sequence** | **Tm** (°C) | **Reference** |
| --- | --- | --- | --- |
| ***tub2*** | | | |
| Zt_tub2_prom_F1 | ATTGCTGTGCTTGTGTTCGA (F) | 55.2 | This work |
| Zt_tub2_prom_R1 | CGACATACCCGTTTGCAAGT (R) | 57.3 | - |
| ***cyp51*** | | | |
| Mg51proF | GTGGCGAGGGCTTGACTAC (F) | 61.0 | Chassot *et al,* 2008 |
| MgCyp51-5'-1r | CAAGTTTCCAGAGGCTGGTC (R) | 59.4 | - |
| ***sdhC*** | | | |
| Zt_sdhC_prom_F1 | GACCGGTCGAATGCTTGC (F) | 58.2 | This work |
| Zt_sdhC_prom_R1 | GGTGAGCTTCTGTGCCAAC (R) | 58.8 | - |
| ***mfs1*** |  |  |  |
| MFS1_consensus_2 | GCAAGGATTCGGACTTGACG (F) | 66.9 | Omrane *et al,* 2017 |
| MFS1_consensus_4 | CTGCCGGTATCGTCGATGAC (R) | 67.8 | ***-*** |

Chassot, C., et al. (2008). Sensitivity of *Cy*p*51* genotypes to DMI fungicides in *Mycosphaerella graminicola*. Modern Fungicides and Antifungals compounds V. G. U. Dehne DW, Kuck KH, Russell PE and Lyr H. Braunschweig, Germany, DPG Seebsterverlag**:** 129-136.

Omrane, S., et al. (2017). "Plasticity of the *MFS1* promoter leads to multidrug resistance in the wheat pathogen *Zymoseptoria tritici*." mSphere **2**(5).

2. **Regular PCR Mix**

| Mix | 1 tube (µL) |
| --- | --- |
| DNA | 1 |
| dNTP (25mM) | 0.24 |
| Primer FW (10 µM) | 0.5 |
| Primer RV (10 µM) | 0.5 |
| Titanium buffer (10X) | 2.5 |
| Titanium *Taq* polymerase (50X) | 0.3 |
| Deionized H_2_O | 19.96 |
| Total volume | 25 |

DNA polymerase: Titanium Taq (Takara Bio)

3. **Regular PCR program**

For *cyp51, sdhC* and *mfs1*

| Initiation | 95°C | 3 min |  |
| --- | --- | --- | --- |
| Denaturation | 95°C | 30 s |  |
| Annealing | Tm°C | 30 s | 35 cycles |
| Elongation | 72°C | 1min |  |
| Final | 72°C | 5 min |  |
|  | 10°C | To the end of the cycle | |

For *tub2*

| Initiation | 95°C | 3 min |  |
| --- | --- | --- | --- |
| Denaturation | 95°C | 30 s | 15cycles |
|  | 68°C | 1mn |  |
| Denaturation | 95°C | 30 s |  |
| Annealing | Tm°C | 30 s | 35 cycles |
| Elongation | 72°C | 1min |  |
| Final | 72°C | 5 min |  |

**Supporting Information Methods S4:** **Quantifying the expression of *tub2*, *cyp51*, *sdhC*, *mfs1* in *Z. tritici* by qRT-PCR**

1. **Origin of tested isolates**

| **Isolate** | **Selection regime** | **Tested genes** |
| --- | --- | --- |
| IPO323 | None (ancestral strain) | *tub2*, *cyp51*, *sdhC* and *mfs1* |
| CS_I_12_A_a | Control with 0.5% ethanol as solvent | *tub2*, *cyp51*, *sdhC* and *mfs1* |
| 0C_I_12_A_a | 0C | *tub2* |
| 0C_II_12_A_b | 0C | *tub2* |
| 0B_I_12_A_b | 0B | *sdhC* |
| 0B_II_12_A_a | 0B | *sdhC* |
| 0P_I_12_A_a | 0P | *cyp51* |
| 0P_II_6_C_b | 0P | *cyp51* |
| 1CP_IV_12_A_a | 1CP | *tub2, cyp51* and *mfs1* |
| 3CP_IV_12_C_b | 3CP | *tub2, cyp51* and *mfs1* |
| 1BC_I_12_C_a | 1BC | *tub2, sdhC* and *mfs1* |
| 2BC_I_12_B_a | 2BC | *tub2, sdhC* and *mfs1* |
| 2BP_I_12_A_a | 2BP | *cyp51, sdhC* and *mfs1* |
| 3BP_IV_12_C_b | 3BP | *cyp51, sdhC* and *mfs1* |
| 1BCP_I_12_C_b | 1BCP | *tub2, cyp51, sdhC* and *mfs1* |

2. **Primers used in qPCR**

| **Primer name** | **Primer sequence (5’-3’)** | **Reference** | **Use** |
| --- | --- | --- | --- |
| β-tubulin_Fw | AACGAGGCTCTCTACGACATCTG | Omrane *et al*, 2017 | Expression of *tub2*, encoding β-tubulin |
| β-tubulin_Rv | GGCGGAGACGAGGTGGTTG |  |  |
| CYP51_Fw | TTCTCTTCCGTGGCAAGTTGTC | S. Patry-Leclaire, unpublished | Expression of *cyp51*, encoding sterol 14α-demethylase |
| CYP51_Rv | GCCGTATGTGATGGTGCTTCC |  |  |
| MFS1_F | CAATGGGGCGGCAGCAAATAC | Omrane *et al*, 2015 | Expression of *mfs1,* encoding a MFS drug transporter |
| MFS1_R | TTGGATGGTGGCGAAGATGAGG |  |  |
| UBC_Fw | GTCTGCGGACCACAATACC | Omrane *et al*, 2017 | Ubiquitine housekeeping gene |
| UBC_RV | CGACCTTTCCTTGCCTCTG |  |  |
| EF1α_Fw | AAGATTGGTGGTATCGGAACAG | Omrane *et al*, 2017 | Elongation factor EF1α housekeeping gene |
| EF1α_Rv | GACTTGACTTCGGTGGTGAC |  |  |
| qPCR_SDHC1_fwd | GGCAACAACTCCTTCCAACAG | Steinhauer *et al,* 2019 | Expression of *sdhC*, encoding the C subunit of the succinate dehydrogenase |
| qPCR_SDHC1_rev | ACCAGGTTATTTGCGGTTTGT |  |  |
| qPCR_SDHC1_probe (CY5/BHQ2) | CCAAGTAACAGCAGCCGCCGTCTCCGAATC |  |  |
| qPCR_TUB1_fwd | AGCGCATGAATGTCTACTTCAA | Steinhauer *et al,* 2019 | β-tubulin housekeeping gene (for *sdhC* only) |
| qPCR_TUB1_rev | CCTTGGCCCAGTTGTTTCC |  |  |
| qPCR_TUB1_probe (HEX / BHQ1) | TCGGTCAGCTCTTCCGCCCAGA |  |  |

*EF1α* and *UBC* were used as reference genes to quantify the expression of *tub2, cyp51* and *mfs1.* *tub2* was used as a reference gene to quantify the expression of *sdhc.*

Steinhauer, D., et al. (2019). "A dispensable paralog of succinate dehydrogenase subunit C mediates standing resistance towards a subclass of SDHI fungicides in *Zymoseptoria tritici*." PLOS Pathogens **15**(12): e1007780.Omrane, S., et al. (2015). "Fungicide efflux and the MgMFS1 transporter contribute to the multidrug resistance phenotype in *Zymoseptoria tritici* field isolates." Environmental Microbiology **17**(8): 2805-2823.

Omrane, S., et al. (2017). "Plasticity of the *MFS1* promoter leads to multidrug resistance in the wheat pathogen *Zymoseptoria tritici*." mSphere **2**(5).

3. **Protocols**

Isolates were grown in the dark at 18°C, with a RH of 70% and shaking at 150 rpm for four days, in 50 mL borosilicate Erlenmeyer flasks containing 25 mL YPD medium and no fungicide. Two independent biological repeats, grown successively, were produced for each isolate. Fungal cells were freeze-dried and RNAs were extracted using a commercial kit (Monarch total RNA Miniprep; New England Biolabs) and the manufacturer’s recommendations. Retrotranscription was achieved using the protoScript II First Strand cDNA Synthesis kit (New England BioLabs) according to the manufacturer’s protocol. qPCRs were achieved as follows, with three technical repeats for each sample:

| Mix | 1 tube (µL) |
| --- | --- |
| cDNA | 2 |
| Primer FW (10 µM) | 0.4 |
| Primer RV (10 µM) | 0.4 |
| Master Mix (2X) | 10 |
| Deionized H_2_O | 7.2 |
| Total volume | 20 |

DNA polymerase: MESA GREEN qPCR MasterMix Plus for SYBR Assay (Eurogentec)

Regular qPCR program for *tub2, cyp51, mfs1, EF1α* and *UBC*

| 95°C | 5 min |  |
| --- | --- | --- |
| 95°C | 15 s | 45 cycles |
| 60°C | 1min |  |

Regular qPCR program for *sdhC*

| 95°C | 3 min |  |
| --- | --- | --- |
| 95°C | 10 s | 45 cycles |
| 60°C | 20 s |  |

4. **Determination of the relative gene expression, 2^-ΔΔCt^**

First, ΔCt values were calculated as:

$$\Delta Ct={Ct}_{i, target gene}-{Ct}_{i, housekeeping gene}$$

with Ct the value of cycle threshold and i each isolate tested.

ΔΔCt values were calculated after the normalization from the control ancestral strain IPO323:

$$\Delta\Delta\mathrm{Ct}= {\Delta\mathrm{Ct}}_{i}-{\Delta\mathrm{Ct}}_{IPO323}$$
